# Standardized pipeline for establishing, expanding, and differentiating airway and alveolar organoids from human BAL fluid

**DOI:** 10.1101/2025.10.13.682184

**Authors:** Taryn B. Gellner, Belinda Chen, Shreyas R. Raini, Mackenzie S. Jackson, Amy K. Kraak, Sophie C. Petta, Margherita Paschini, Lynn M. Schnapp, Carla F. Kim, Monica Yun Liu

**Affiliations:** Division of Allergy, Pulmonary, and Critical Care, Department of Medicine, University of Wisconsin-Madison, Wisconsin, USA; Stem Cell Program, Division of Hematology/Oncology and Pulmonary & Respiratory Diseases, Department of Pediatrics, Boston Children’s Hospital, Massachusetts, USA; Department of Genetics, Harvard Medical School, Massachusetts, USA

## Abstract

Lung organoids are versatile experimental models, but their broader use in studying human disease is limited by the scarcity of starting material and the complexity of current methods. To align organoid technology with common clinical practice, we developed airway and alveolar organoids using cells obtained from patients’ bronchoalveolar lavage (BAL) fluid. Building on existing techniques, we showed that BAL is a reliable, accessible source of primary human epithelial cells, yielding airway and alveolar organoids within 10 days. Organoids can then be expanded over many passages for downstream analysis. Our streamlined methods do not require cell sorting or other complex procedures, all cells are derived from a single patient, and media are based on serum-free, chemically-defined formulations. Here, we present detailed protocols for organoid establishment, standardized passaging and phenotyping, and differentiation of both airway and alveolar models. We provide a time course of BAL-derived airway organoid differentiation at air-liquid interface, and we demonstrate proof of principle for differentiation of BAL-derived alveolar organoids in 3D culture. These methods can be readily adapted to generate and characterize organoids from lung tissue, tracheobronchial specimens, or other primary cells from humans or mice, expanding the potential to use lung organoids for disease modeling.

**NEW AND NOTEWORTHY:** We provide streamlined protocols to generate both airway and alveolar epithelial organoids from a single, clinical BAL specimen. From standardized specimen collection to organoid plating to passaging and differentiation, we show that rare, primary epithelial cells in BAL can give rise to all the major airway and alveolar cell types. Our serum-free, feeder-free, sorting-free methods offer a simplified starting point for using patient-derived organoids to model lung disease.

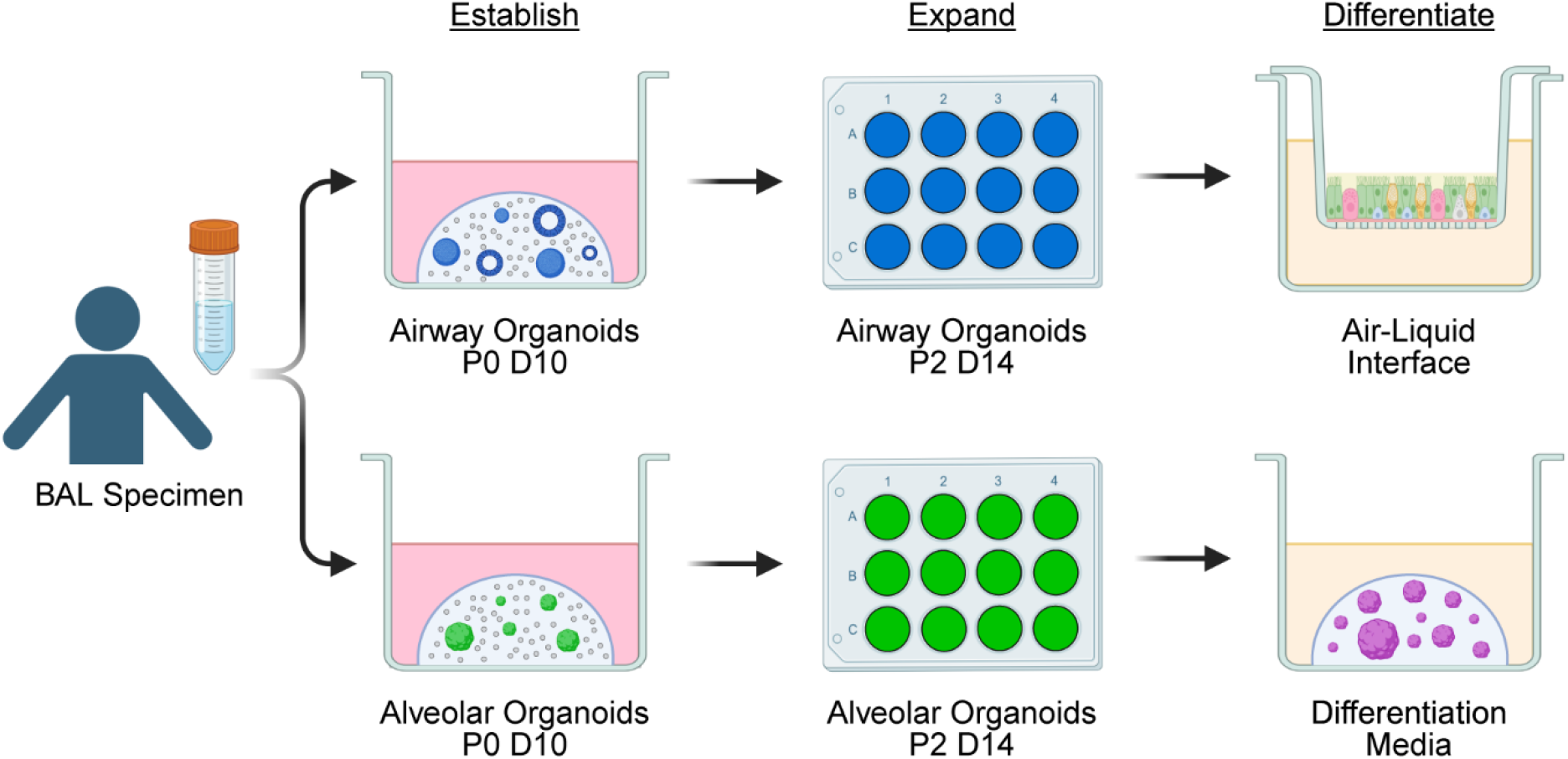
Graphical Abstract. Standardized pipeline for BAL organoids. Created in BioRender.

## INTRODUCTION

Organoids are three-dimensional (3D), self-assembled groups of cells that mimic a particular tissue, incorporating multiple cell types and often recapitulating form and function in ways that traditional 2D culture cannot.^1^ Human lung organoids vary widely in their starting material, cellular composition, culture format, media ingredients, and matrix material.^2^ Here, we focus on human adult organoids from primary cells that are grown in conditions that favor either airway or alveolar epithelial cell expansion. Airway organoids were first generated from basal stem cells in primary bronchial epithelial tissue.^3^ Alveolar organoids were first established using alveolar type 2 (AT2) stem cells from lung tissue, requiring MRC5 cells (a fetal human lung fibroblast cell line) for support.^4^ These and subsequent systems opened new windows into human biology and disease, but there were key limitations. Cultures generally survived only a few passages. The media contained animal serum, which has unknown and variable components that can change cellular identity. The procedures were labor-intensive, calling for lung harvesting, digestion, and cell sorting on the same day, while supporting cells had to be grown separately and harvested in time to co-culture with epithelial organoids.

Long-term expanding airway organoids were developed in 2019,^5^ and alveolar organoids followed in 2020,^6,7^ showing that primary human stem cells could be maintained for months *in vitro* without losing their identity. These newer methods used defined chemicals and growth factors to modulate specific pathways known to drive lung stem cell development, maintenance, and differentiation, supplanting the role of supporting cells. These “serum-free, feeder-free” approaches made cultures more reproducible and simplified the organoid workflow. Nonetheless, most methods required sorting cells from lung tissue, which is difficult to obtain and not available for most disease states. Organoids derived from induced pluripotent stem cells (iPSCs) provided a more flexible cell source but require complex manipulation.^8–11^ A major advance came with showing that culture media alone is sufficient to select for airway cell outgrowth, without cell sorting. It became possible to establish airway organoids from bronchoalveolar lavage (BAL) fluid,^5^ tracheal aspirates,^12^ and nasal brushings^13^—accessible sources of primary cells that contain only rare stem cells proficient at organoid formation. Our group further simplified and characterized airway organoids from human BAL fluid and established alveolar organoids from BAL as well.^14^ Combining serum-free, feeder-free, sorting-free approaches, we can now progress from BAL collection to organoid plating in approximately 4 hours using standard laboratory equipment and techniques. Here, we present updated protocols for culturing airway and alveolar BAL organoids that further standardize sample collection, passaging, and phenotyping. We also demonstrate examples of differentiating airway and alveolar stem cells in 2D and 3D formats, respectively, showing that BAL organoids remain capable of recapitulating a full array of relevant cell types for disease modeling and advanced applications.

## MATERIALS AND METHODS

### Materials

All materials and catalog numbers are listed in **Table 1**, along with antibody dilutions. We prepare stock solutions according to the manufacturers’ instructions, and we reconstitute proteins in Dulbecco’s phosphate-buffered saline (DPBS) with 0.1% (w/v) bovine serum albumin (BSA) as a stabilizing agent. We store small molecules at -20°C and proteins at -80°C and use within two freeze-thaw cycles. We use Matrigel that is optimized for organoid culture (Corning 356255). Once thawed, we store Matrigel at 4°C and use within 2 weeks.

**Table 1.** Complete list of materials and catalog numbers, with antibody dilutions.

| Reagent or Resource | Source | Catalog Number |
| --- | --- | --- |
| **Cell Culture Media Components** |  |  |
| Advanced DMEM/F-12 | Thermo Fisher | 12634010 |
| GlutaMAX Supplement | Thermo Fisher | 35050061 |
| HEPES (1X) | Thermo Fisher | 15630080 |
| Primocin | InvivoGen | ant-pm-1 |
| B-27 Supplement (50X) | Thermo Fisher | 17504044 |
| N-2 Supplement (100X) | Thermo Fisher | 17502048 |
| Insulin-Transferrin-Selenium (100X) | Thermo Fisher | 41400045 |
| Nicotinamide | Millipore Sigma | N0636 |
| N-Acetyl-L-cysteine | Millipore Sigma | A9165 |
| Heparin | Millipore Sigma | H3149 |
| Recombinant Human EGF | Thermo Fisher | AF-100-15 |
| Recombinant Human R-Spondin-1 | Thermo Fisher | 120-38 |
| Recombinant Human Noggin | Thermo Fisher | 120-10C |
| Recombinant Human FGF-10 | Thermo Fisher | 100-26 |
| Recombinant Human FGF-7 (KGF) | Thermo Fisher | 100-19 |
| Recombinant Human Heregulinβ-1 | Thermo Fisher | 100-03 |
| Recombinant Human IL-1β | Thermo Fisher | 200-01B |
| CHIR 99021 | Tocris | 4423 |
| A 83-01 | Millipore Sigma | SML0788 |
| SB 202190 | Tocris | 1264 |
| SB 431542 | Tocris | 1614 |
| BIRB 796 | Tocris | 5989 |
| ROCK Inhibitor Y27632 | ATCC | ACS-3030 |
| Amphotericin B | Millipore Sigma | A2942 |
| Gentamicin | Thermo Fisher | 15710064 |
| PneumaCult-Ex Plus Medium | StemCell Technologies | 05040 |
| PneumaCult-ALI Medium | StemCell Technologies | 05001 |
| PneumaCult-ALI-S Medium | StemCell Technologies | 05050 |
| Hydrocortisone Stock Solution | StemCell Technologies | 07925 |
| Human Serum | Millipore Sigma | H4522 |
| IMDM | Thermo Fisher | 12440053 |
| Ham's F-12 | Corning | 10-080-CV |
| Bovine Serum Albumin (7.5% w/v solution) | Thermo Fisher | 15260037 |
| L-Ascorbic Acid | Millipore Sigma | A4544 |
| 1-Thioglycerol (MTG) | Millipore Sigma | M6145 |
| TRULI (LATS1/2 Inhibitor) | MedChem Express | HY-138489 |
| Dexamethasone | Millipore Sigma | D4902 |
| 8-Bromoadenosine 3',5'-Cyclic Monophosphate Sodium Salt (cAMP) | Millipore Sigma | B7880 |
| 3-Isobutyl-1-Methylxanthine (IBMX) | Millipore Sigma | I5879 |
| **Antibodies and Dyes** |  |  |
| Chicken Polyclonal IgY anti-Human KRT5 (1:500) | BioLegend | 905901 |
| Goat Monoclonal IgG anti-Human KRT5 (1:500) | Abcam | ab321868 |
| Goat Polyclonal IgG anti-Human KRT13 (1:200) | Abcam | ab79279 |
| Rabbit Monoclonal IgG anti-Human Acetyl-α-Tubulin (1:200) | Cell Signaling Technology | 5335S |
| Mouse Monoclonal IgG1k anti-Human SCGB1A1 (1:200) | Santa Cruz Biotechnology | sc-365992 |
| Rabbit Polyclonal IgG anti-Human SCGB1A1 (1:200) | Thermo Fisher | 26909-1-AP |
| Goat Polyclonal IgG anti-Human SCGB3A2 (1:200) | R&D Systems | AF3545 |
| Mouse Monoclonal IgG1k anti-Human MUC5AC (1:200) | Thermo Fisher | MA5-12178 |
| Rabbit Polyclonal IgG anti-Human pro-SPC (1:500) | Seven Hills Bioreagents | WRAB-9337 |
| Goat IgG anti-Human AGER (1:200) | R&D Systems | AF1145 |
| Donkey anti-Chicken IgY Highly Cross Adsorbed, Alexa Fluor 647 (1:500) | Thermo Fisher | A78952 |
| Donkey anti-Rabbit IgG Highly Cross-Adsorbed, Alexa Fluor Plus 488 (1:500) | Thermo Fisher | A32790 |
| Donkey anti-Mouse IgG Highly Cross-Adsorbed, Alexa Fluor Plus 594 (1:500) | Thermo Fisher | A32744 |
| Donkey anti-Mouse IgG Highly Cross-Adsorbed, Alexa Fluor Plus 647 (1:500) | Thermo Fisher | A32787 |
| Donkey anti-Goat IgG Highly Cross-Adsorbed, Alexa Fluor Plus 594 (1:500) | Thermo Fisher | A32758 |
| Phalloidin, Alexa Fluor Plus 405 | Thermo Fisher | A30104 |
| DAPI | Millipore Sigma | D9542 |
| PE anti-Human CD45 Antibody (1:100) | BioLegend | 304008 |
| PE/Cyanine7 anti-Human CD326 (EpCAM) Antibody (1:100) | BioLegend | 324222 |
| APC anti-Human CD271 (NGFR) Antibody (1:100) | BioLegend | 345108 |
| Mouse IgM anti-Human HT2-280 (1:50) | Terrace Biotech | TB-27AHT2-280 |
| Goat anti-Mouse IgM Cross Adsorbed Secondary Antibody, DyLight 488 (1:200) | Thermo Fisher | SA5-10150 |
| UltraComp eBeads Compensation Beads | Thermo Fisher | 01-2222 |
| **Other Reagents** |  |  |
| Matrigel Matrix for Organoid Culture, Phenol Red-free, LDEV-free | Corning | 356255 |
| Liberase TM | Millipore Sigma | 5401127001 |
| Deoxyribonuclease I (DNase) | Millipore Sigma | D4527 |
| TrypLE Express Enzyme (1X), No Phenol Red | Thermo Fisher | 12604021 |
| Bambanker Cell Freezing Medium | Bulldog Bio | BB05 |
| Cryo-SFM Plus Freezing Medium | PromoCell | C-29922 |
| MycoFluor Mycoplasma Detection Kit | Thermo Fisher | M7006 |
| Trypan Blue Solution, 0.4% | Thermo Fisher | 15250061 |
| DPBS, 1X without Calcium and Magnesium | Corning | 21-031-CV |
| EDTA | Millipore Sigma | E6758 |
| Dimethyl Sulfoxide | Millipore Sigma | 472301 |
| Neutral Buffered Formalin, 10% | Fisher Scientific | STL286001 |
| Triton X-100 | Millipore Sigma | T8787 |
| Normal Donkey Serum | Jackson ImmunoResearch | 017-000-121 |
| ProLong Gold Antifade Mountant | Thermo Fisher | P36930 |
| **Other Materials** |  |  |
| 12-well Clear Flat Bottom TC-treated Plate, Sterile | Corning | 353043 |
| 24-well Clear Flat Bottom TC-treated Plate, Sterile | Corning | 353047 |
| 48-well Clear Flat Bottom TC-treated Plate, Sterile | Corning | 353078 |
| 6.5 mm Transwell with 0.4 μm Pore Polyester Membrane Insert, Sterile | Corning | 3470 |
| Light Sensitive Amber 50 ml Centrifuge Tubes | VWR | 89079-535 |
| 2-Chip Disposable Hemocytometer | Bulldog Bio | DHC-N51 |
| 2 ml External Threaded Polypropylene Cryogenic Vial, Self-Standing with Round Bottom | Corning | 430659 |
| CoolCell LX, Cell Freezing Container, for 12 x 1 mL or 2 mL Cryogenic Vials | Corning | 432002 |
| Greiner Bio-One Cell Strainer, 70 μm, for 1.5 to 15 ml tubes | Fisher Scientific | 07-001-107 |
| Greiner Bio-One Cell Strainer, 40 μm, for 1.5 to 15 ml tubes | Fisher Scientific | 07-001-106 |
| Superfrost Plus Microscope slides | Fisher Scientific | 1255015 |
| Micro Cover Glasses, 22x22 mm, No. 1.5 | Fisher Scientific | 12541016 |
| Super HT PAP Hydrophobic Slide Marker | Research Products International | 195506 |
| 5 mL Round Bottom Polystyrene Flow Cytometry Tube, with Cell Strainer Snap Cap | Corning | 352235 |
| **Software** |  |  |
| GraphPad Prism | https://www.graphpad.com/ |  |
| Fiji | https://imagej.net/software/fiji/ |  |
| Flow Jo | https://www.flowjo.com/ |  |

To develop these protocols, we used BAL and lung tissue from a variety of sources. Most specimens were from lung transplant recipients at Mass General Brigham in Boston or the University of Wisconsin-Madison because these patients undergo frequent bronchoscopies. Control specimens included BAL from healthy volunteers and lung tissue from deceased donors whose organs could not be used for transplant. When we obtained specimens from other repositories, they were deidentified, so the work was classified as non-human subjects research and was exempted from Institutional Review Board approval. We now also run a biorepository for lung transplant recipients at UW-Madison, which is approved under IRB #2023-1445. In all cases, specimens were originally obtained with written informed consent from all donors or their legally authorized representative.

### Media

We previously developed our organoid media based on published formulations, adding our own slight variations.^5,6,14,15^ For simplicity, we denote the media as Airway or Alveolar, and we streamlined the Base Media to be the same for both (**Table 2**). We also add HRG1-β1 to Airway Media^14^ and IL-1β to Alveolar Media^6,14,15^ for the first four days after initial plating (Passage 0) to promote cell survival.

**Table 2.** Media recipes.

| Reagent | Stock Concentration | Final Concentration | Volume for 50 ml |
| --- | --- | --- | --- |
| **Base Media** |  |  |  |
| Advanced DMEM/F12 |  |  | 47.9 ml |
| GlutaMAX | 100X | 1X | 500 µl |
| HEPES | 1 M | 10 mM | 500 µl |
| Primocin | 50 mg/ml | 100 µg/ml | 100 µl |
| B-27 | 50X | 1X | 1 ml |
| **BAL Buffer** |  |  |  |
| Base Media |  |  | 49.7 ml |
| Amphotericin B | 250 µg/ml | 250 ng/ml | 50 µl |
| Gentamicin | 10 mg/ml | 50 µg/ml | 250 µl |
| ROCK Inhibitor Y27632 | 10 mM | 5 µM | 25 µl |
| **Passaging Buffer** |  |  |  |
| Base Media |  |  | 50 ml |
| ROCK Inhibitor Y27632 | 10 mM | 5 µM | 25 µl |
| **Airway Media** |  |  |  |
| Base Media |  |  | 49.5 ml |
| Nicotinamide | 1 M | 5 mM | 250 µl |
| N-Acetyl-L-cysteine | 1.25 M | 1.25 mM | 50 µl |
| Recombinant Human R-Spondin-1 | 500 µg/ml | 500 ng/ml | 50 µl |
| Recombinant Human Noggin | 100 µg/ml | 100 ng/ml | 50 µl |
| Recombinant Human FGF-10 | 100 µg/ml | 100 ng/ml | 50 µl |
| Recombinant Human FGF-7 | 100 µg/ml | 25 ng/ml | 12.5 µl |
| A 83-01 | 5 mM | 500 nM | 5 µl |
| SB 202190 | 5 mM | 500 nM | 5 µl |
| ROCK Inhibitor Y27632 | 10 mM | 5 µM | 25 µl |
| Recombinant Human Heregulinβ-1 | 50 µg/ml | 50 ng/ml | 1 µl per ml Airway Media, first 4 days only |
| **Alveolar Media** |  |  |  |
| Base Media |  |  | 48.9 ml |
| N-2 | 100X | 1X | 500 µl |
| Insulin-Transferrin-Selenium | 100X | 1X | 500 µl |
| N-Acetyl-L-cysteine | 1.25 M | 1.25 mM | 50 µl |
| Heparin | 50 µg/µl | 5 µg/ml | 5 µl |
| Recombinant Human FGF-10 | 100 µg/ml | 10 ng/ml | 5 µl |
| Recombinant Human EGF | 1 µg/µl | 50 ng/ml | 2.5 µl |
| CHIR 99021 | 10 mM | 3 µM | 15 µl |
| SB 431542 | 50 mM | 10 µM | 10 µl |
| BIRB 796 | 10 mM | 1 µM | 5 µl |
| ROCK Inhibitor Y27632 | 10 mM | 5 µM | 25 µl |
| Recombinant Human IL-1β | 10 μg/ml | 10 ng/ml | 1 μl per ml Alveolar Media, first 4 days only |
| **Alveolar Differentiation Media (ADM, from ref. 6)** |  |  |  |
| Base Media |  |  | 45 ml |
| N-Acetyl-L-cysteine | 1.25 M | 1.25 mM | 50 μl |
| Heparin | 50 μg/μl | 5 μg/ml | 5 μl |
| Recombinant Human EGF | 0.1 μg/μl | 5 ng/ml | 2.5 μl |
| Recombinant Human FGF-10 | 10 μg/ml | 1 ng/ml | 5 μl |
| Human Serum |  | 10% (v/v) | 5 ml |
| **L-DCI Differentiation Media (from ref. 22)** |  |  |  |
| IMDM |  |  | 37.5 ml |
| Ham's F12 |  |  | 12.5 ml |
| GlutaMAX | 100X | 1X | 500 μl |
| Primocin | 50 mg/ml | 100 μg/ml | 100 μl |
| B-27 | 50X | 0.5X | 500 μl |
| N-2 | 100X | 0.5X | 250 μl |
| Bovine Serum Albumin | 7.5% (w/v) | 0.05% (w/v) | 333 μl |
| L-Ascorbic Acid | 50 mg/ml | 50 μg/ml | 50 μl |
| 1-Thioglycerol (MTG) | 150 mM | 450 μM | 150 μl |
| TRULI (LATS1/2 Inhibitor) | 50 mM | 10 μM | 10 μl |
| Dexamethasone | 1.25 mM | 50 nM | 2 μl |
| 8-Bromo-cAMP | 50 mM | 100 μM | 100 μl |
| 3-Isobutyl-1-Methylxanthine (IBMX) | 100 mM | 100 μM | 50 μl |

We use Base Media within 6 weeks, BAL and Passaging Buffer within 4 weeks, and complete Airway and Alveolar Media within 2 weeks.

### Collecting BAL

We prefer to use fresh, excess BAL from clinical bronchoscopies. We recommend standardizing the BAL procedure. UW pulmonary clinicians follow guidelines from the International Society of Heart and Lung Transplantation (ISHLT), which call for instillation of two 50-ml aliquots of saline and pooling the return.^16^ A similar alternative is three 40-ml aliquots. We do not add additional volume for research purposes. We reserve the first 35 ml of BAL return for clinical lab testing, then use the excess for our studies. We find that 5 ml works well to establish organoids (minimum 1 ml), but more volume is helpful for additional technical replicates or biobanking. We transport the specimen on ice and use or freeze the cell pellet within 4 hr.

### Establishing Organoid Cultures

Here, we provide a step-by-step protocol for establishing organoid cultures from fresh BAL fluid. Our methods are derived primarily from Sachs et al.^5^ but are further simplified. For example, we use enzymatic digestions that avoid passing cells through flamed Pasteur pipettes, and we use completely serum-free methods. We were the first group to establish BAL-derived alveolar organoid cultures using the same protocol. Our methods do not require feeder cells or cell sorting, which significantly reduces processing time and stress on the cells; this also means that our epithelial cell monocultures contain only cells from a single patient. Having now replicated this technique on over 60 biological specimens, plus additional controls, we note tips and troubleshooting strategies that have allowed for relatively easy uptake of these methods. Our full pipeline is readily scalable and adaptable; we briefly comment on protocol variations at the end of each section.

All steps are performed in a biosafety cabinet with sterile technique.

- **Note:** Because media is light-sensitive, we work with the hood light *off*, relying on indirect lighting from the room.
- **Important:** For infection control, we use BAL specimens from biorepositories that collect limited clinical data, so that specimens can be screened for viral infections or drug-resistant organisms. We also include antimicrobials in our media and test all cells for Mycoplasma at least once monthly.

A) Approximate time for establishing organoid cultures: 3-4 hr
  A1) Pre-warm 12-well tissue culture plates to 37°C.
  A2) In a conical tube, spin fresh BAL fluid at 800*g* for 10 min at 4°C. This speed helps to collect all cells, including small mucus plugs.
    - *Optional:* Save an aliquot of whole BAL fluid and/or supernatant for biobanking.
  A3) Aspirate the supernatant and resuspend the cell pellet in a volume of BAL Buffer appropriate for counting, usually 1 ml. Count live and total cells (e.g. using Trypan Blue) and note the percent viability.
  A4) Transfer up to 1 million live cells to a 15-ml conical. Bring the volume to 5 ml of BAL Buffer.
  Freeze any excess cells (see “Establishing Organoids: Protocol Variations” below).
    - *Example:* If 3,200,000 live cells were counted in 1 ml BAL Buffer at step **A3**, transfer 313 µl (1,000,000 cells) to a 15-ml conical for plating. Freeze the remaining cells.
  A5) To the BAL cells in 5 ml BAL Buffer, add Liberase TM (1:100, final concentration 50 µg/ml) and DNase I (1:500, final concentration 20 U/ml). This enzymatic digestion breaks up clumps of cells and mucus to generate a single-cell suspension.
  A6) Gently agitate at 37°C for 20 min. If needed, manually mix by inverting every 10 min to ensure thorough digestion.
  A7) Place a a sterile 70-µm mesh filter inside a clean 15-ml conical tube. Pre-wet the filter with 1 ml BAL Buffer.
  A8) Pass digested cells through the 70-µm mesh filter into the clean conical tube. Use an additional 1 ml BAL Buffer to wash the filter and maximize yield.
  A9) *Perform remaining steps on ice or at 4°C*.
  A10) Spin at 200*g* for 10 min. This speed is optimized to collect live cells post-digestion while eliminating dead cells and debris.
  A11) The BAL cell pellet should appear white. If the cell pellet is red, add the following red blood cell (RBC) lysis step: resuspend the cell pellet in 1 ml RBC Lysis Buffer (150 mM ammonium chloride, 10 mM potassium bicarbonate, and 0.1 mM EDTA). Incubate at room temperature for *no more than 2 min*. Add 5 ml BAL Buffer to stop the lysis and spin at 200*g* for 10 min.
    - **Note:** BAL cells can also appear gray or brown depending on the patient (e.g. with smoking). We do not use RBC lysis for these samples, since the pigmentation usually comes from macrophages.
  A12) Resuspend the cell pellet in 5 ml BAL Buffer (wash #1). Do not use less than 5 ml, since it is important to remove any remaining enzymes and debris. Spin at 200*g* for 10 min.
  A13) Repeat for another wash (wash #2).
  A14) Resuspend cell pellet in a volume of BAL Buffer appropriate for counting, usually 1 ml. Count live and total cells and note the percent viability.
    - **Note:** The post-digestion count often differs from the pre-digestion count and may be higher due to cells freed from mucus clumps. Viability is also improved by washing and should be at least 90%.
    - *Optional:* To perform cytology staining on BAL cells, set aside 100,000 live cells in 400 µl media with 10% FBS for Cytospin. This is enough to make two slides of 50,000 cells each. We spin at 300 rpm for 3 min using a Cytospin 4 Centrifuge, then immediately fix the cells in 10% formalin for 10 min.
  A15) Spin at 200*g* for 10 min.
  A16) Remove as much supernatant as possible without disturbing the cell pellet. Cells are now ready for plating. *For Passage 0, each well will have 100,000 ± 10,000 live cells*. Having started with no more than 1 million cells at step **A4** (pre-digestion), and re-counting at step **A14** (post-digestion), we would expect to be plating no more than 12 wells. Half the wells will receive Airway Media, while the other half will receive Alveolar Media, so that the same starting population of cells will undergo selection for different cell types.
  A17) Carefully resuspend ∼100,000 live cells per 50 µl Matrigel.
  *Example:* If there are 930,000 live cells, resuspend in 450 µl Matrigel, which will be enough to plate 9 wells (e.g. 4 Airway, 5 Alveolar).
    - **Important:** Since Matrigel solidifies at temperatures above 10°C, *keep all materials cold/on ice*. Handle tubes containing Matrigel by the tops only to avoid exposing Matrigel to warm fingers. Hold the pipette tips against the cold plastic on the inside of the tube for 10 seconds before touching the Matrigel. The tube should stay in contact with ice at all times.
    - **Important:** Pipette slowly, since Matrigel is viscous, and pause for 3 sec to allow the full volume of Matrigel to fill the pipette. Mix the cells evenly and *avoid bubbles*.
  A18) Plate 50-µl drops in the center of each well of the pre-warmed 12-well plate (**Figure 1A**). Pipette the Matrigel up and down between plating each drop to ensure an even cell mixture. At the end, distribute any remaining Matrigel into the wells. Carefully remove any bubbles.
    - *Tip:* Put the plate on top of a plastic microfuge tube rack so that it cools less quickly than on the metal surface of the biosafety cabinet. Pipetting Matrigel onto the pre-warmed plate helps to establish a stable drop.
    - *Tip:* Reverse pipetting can help to minimize bubbles. Depress the pipette plunger past the first stop to pick up a little extra Matrigel. Then, go up and down to the first stop to dispense 50-µl aliquots.
    - *Troubleshooting:* If drops consistently run to the side of the well, instead of staying domed in the middle, we recommend checking the following: make sure the surface is level; check that Matrigel is less than 2 weeks old and has a protein concentration of at least 8 mg/ml; at step **A16**, remove as much supernatant as possible from the cell pellet to avoid diluting the Matrigel; finally, consider plating smaller drops (e.g. 40 µl at the same cell density).
  A19) Carefully transport the plate to the incubator. Let drops solidify for 25 min at 37°C.
  A20) While waiting, warm up 1 ml of Airway or Alveolar Media per well to 37°C.
    - **Important:** Add human HRG1-β1 (1:1,000, final concentration 50 ng/ml) to Airway Media and human IL-1β (1:1,000, final concentration 10 ng/ml) to Alveolar Media.
    - **Note:** HRG1-β1 and IL-1β are for the initial plating only, when organoids are first established. They are not used for subsequent passages.
  A21) Once Matrigel has solidified, gently add 1 ml of Airway or Alveolar Media down the side of each well.
  A22) This is Passage 0 Day 0 (P0 D0). Incubate cells at 37°C, 5% CO2.
  A23) Change media on Day 4 to remove HRG1-β1 and IL-1β and feed again every 3-4 days thereafter.
  A24) Count and image airway and alveolar organoids at Passage 0 Day 10 (P0 D10). We quantify organoid forming efficiency (OFE) for each condition as the total number of organoids divided by the number of live cells plated.

**Figure 1.**
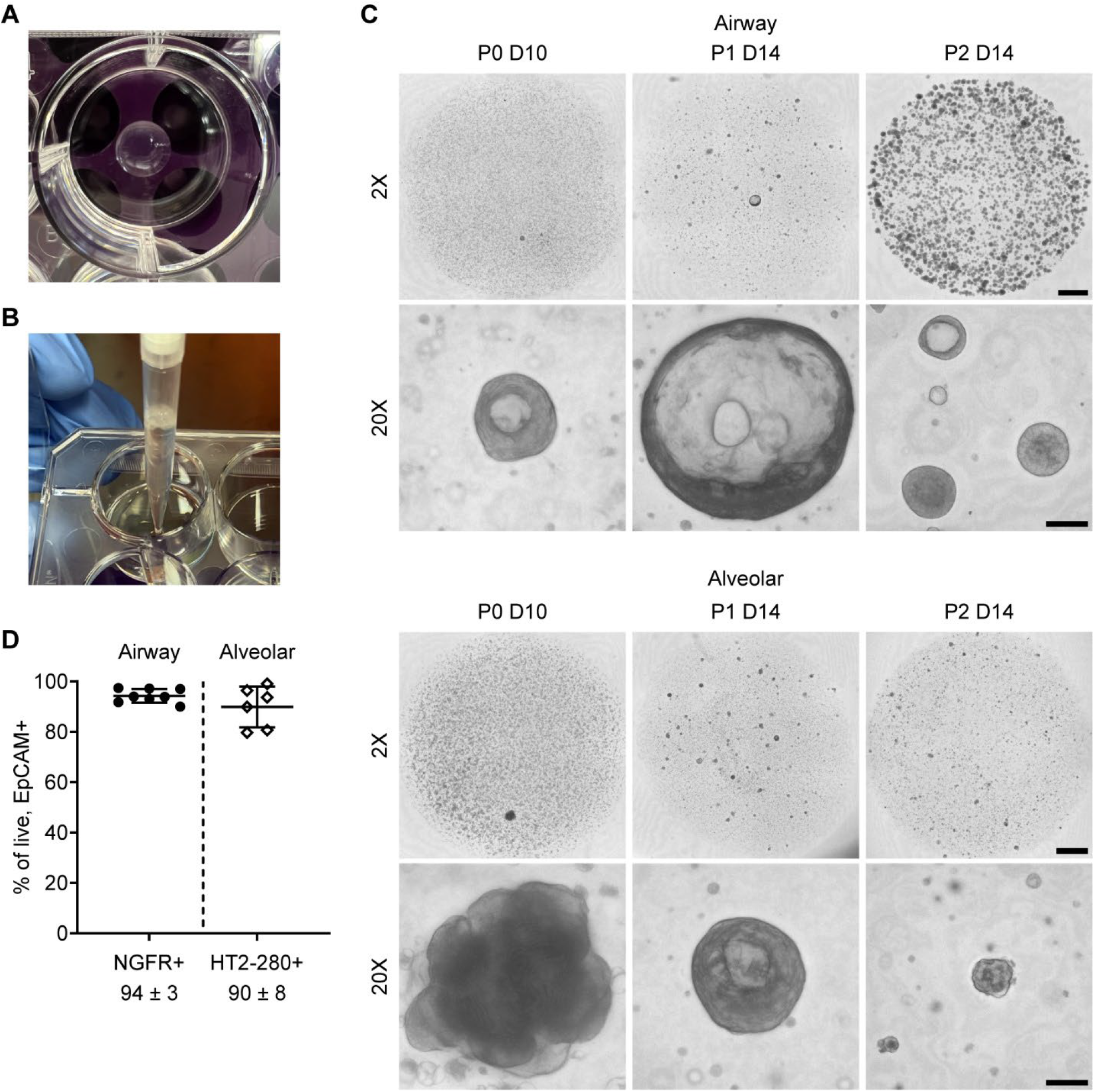
Key steps during organoid culture and expected results. **(A)** Example of cells plated in a 50-µl drop of Matrigel on a pre-warmed 12-well plate. Once solidified, the Matrigel drop can be covered with Airway or Alveolar Media. **(B)** To passage organoids, after incubation in TrypLE, organoids are mechanically sheared through a pipette tip pressed perpendicularly against the bottom of the plate. **(C)** Example of BAL-derived airway (top) and alveolar (bottom) organoids at Passage 0 Day 10, Passage 1 Day 14, and Passage 2 Day 14. Organoid morphologies are variable at P0-P1 and become more uniform from P2 onward. Brightfield images were acquired on a Keyence BZ-X810 microscope. A z-stack was taken covering the entire Matrigel drop (at 2X magnification) or examples of organoids (at 20X magnification), and the Keyence “full focus” feature was used to flatten the stack into one 2D image. Scale bars are 1,000 µm at 2X and 100 µm at 20X. **(D)** Flow cytometry of airway and alveolar organoid cultures at Passage 2 Day 14-18 shows that the culture media successfully select for either NGFR+ airway basal cells or HT2-280+ alveolar type 2 cells. After gating for live, single cells, >95% of cells are EpCAM+ epithelial cells. The percentage of epithelial cells that are NGFR+ in airway cultures or HT2-280+ in alveolar cultures is given as mean ± SD. Updated from ref. 14.

#### Establishing Organoids: Protocol Variations

- Instead of plating all cells fresh, specimens can be cryopreserved (pre-digestion). Spin the BAL specimen at 800*g* for 10 min at 4°C. Resuspend cell pellets in CryoSFM or Bambanker serum-free freezing medium, up to 1 million cells per 1 ml, and transfer to cryovials. Cool the vials slowly at -80°C for at least 24 hr, then transfer cells to liquid nitrogen. We have found that cell viability is better preserved with CryoSFM or Bambanker, though 90% fetal bovine serum (FBS) with 10% DMSO can be used as well.
  - To thaw cells, for each cryovial containing 1 ml (up to 1 million cells), transfer the cell suspension into 5 ml BAL Buffer. Spin at 800*g* for 10 min at 4°C, wash with another 5 ml BAL Buffer, spin, and resuspend in 5 ml BAL Buffer for digestion. Proceed with step **A5**.

- Lung tissue can be used instead of BAL fluid to establish both airway and alveolar organoid cultures, without the need for cell sorting. We start with approximately 1 cm^3^ of minced lung tissue, chopped using sterile scissors or a razor blade so that the pieces fit through a 5-ml serological pipette tip. Resuspend the minced tissue in 8 ml BAL Buffer in a 15-ml conical tube. Digest with Liberase TM and DNase at the concentrations above for 1 hr at 37°C. Proceed with step **A7**.

### Passaging Organoid Cultures

We perform the first passage at P0 D10 and then every 14 days thereafter. The earlier first passage seems to capture the newly-established organoids in their active growth phase, helping to promote further expansion. In our initial publication, we described passaging at 1:1 to 1:4 ratios without cell counting.^14^ We have since standardized our protocol to make it more quantitative. Here, we describe our updated protocol, including a longer dissociation time to obtain a single-cell suspension. The cells are then counted, allowing for measurement of organoid forming efficiency (OFE) and organoid growth rate at each passage.

- **Important:** Every time cells are exposed to TrypLE is one passage.
- **Important:** Each BAL specimen has now been grown in two separate conditions: Airway and Alveolar. Both sets of cells are passaged on the same day, using the same protocol, in separate tubes. To avoid redundancy, we will not state this at every protocol step.

B) Approximate time for passaging organoid cultures: 3 hr
  B1) Pre-warm new 12-well tissue culture plates to 37°C.
  B2) Aspirate media from the wells. To dissociate the Matrigel and organoids, use 1 ml of TrypLE per 2 wells. Add 1 ml TrypLE to the first well. While ejecting, use the pipette tip to scrape Matrigel gently off the bottom of the plate. Pipette up and down one time to break up Matrigel. Pipette up all the Matrigel pieces and transfer into a second well, scraping apart the Matrigel in the second well. Pipette up and down one more time.
    - **Note:** If desired, every well can be dissociated with 1 ml TrypLE. We combine two wells into one to save on the volume of media, so that all steps can be performed in a 15-ml conical tube.
  B3) Incubate the plate at 37°C for 10 min.
  B4) While cells are incubating, prepare a 15-ml conical tube with enough Passaging Buffer to dilute TrypLE by 5-fold. Place on ice.
    - *Example:* For 5 wells of organoids, we use 3 ml TrypLE and add 12 ml Passaging Buffer.
  B5) Pipette up the organoids and shear them by ejecting while pressing the pipette tip perpendicularly against the bottom of the plate (**Figure 1B**). There should be mild resistance while ejecting, and the organoid suspension should squeeze steadily through the space between the pipette tip and the plate. Shear two times. Minimize bubbles.
  B6) Check digestion under a microscope at low magnification (e.g. 4X objective). Initially, organoids will still be largely intact, but the Matrigel should be dissolved. Incubate for another 10 min.
  B7) Repeat steps **B5**-**B6** until the organoids are mostly single cells, up to 30-40 min total.
    - **Note:** Stop if the cells form strands that don’t break up with pipetting, as this indicates they are dying and clumping up. Normally, this should not happen.
  B8) Transfer cells into the prepared tube of Passaging Buffer to inactivate TrypLE. Use buffer to wash the wells to recover as many cells as possible.
  B9) Filter cells through a 40-µm mesh into a new conical tube. Wash the filter as before to maximize cell recovery.
    - *Exception:* If there were <10 organoids to begin with, skip filtering so as not to lose precious cells.
  B10) Spin at 200*g* for 10 min.
  B11) Resuspend cell pellet in a volume of Passaging Buffer appropriate for counting. Count live cells.
  B12) Spin at 200*g* for 10 min.
  B13) Remove as much supernatant as possible without disturbing the cell pellet. Cells are now ready for plating. *For Passage 1 onward, each well will have 20,000 live cells*. We plate up to 6 wells each in Airway and Alveolar conditions for P1, up to 12 wells each for P2, and up to 3 wells each for higher passages. Freeze any excess cells.
    - **Note:** The cell count includes all types of cells, but under the selective media conditions, immune cells are eliminated while epithelial cells proliferate. The rate of epithelial cell expansion varies by specimen, so it is possible to have fewer wells in P1 than in P0.
  B14) As described in steps **A17**-**A21**, carefully resuspend 20,000 live cells per 50 µl Matrigel. Plate 50-µl drops in the pre-warmed 12-well plate. Let drops solidify for 25 min at 37°C and add 1 ml per well of pre-warmed Airway or Alveolar Media (without HRG1-β1 and IL-1β).
  B15) This is Passage 1 Day 0. Incubate cells at 37°C, 5% CO2. Feed with fresh media every 3-4 days.
  B16) Count and image airway and alveolar organoids at Day 14 of every passage.

By Passage 2 Day 14, both airway and alveolar organoids should have reached maturity and expanded enough for desired applications (**Figure 1C**). They can be differentiated, harvested for flow cytometry or “-omics” analysis,^14^ fixed, lysed for DNA/RNA isolation, passaged further, cryopreserved, etc. (Logistically, we tend to passage at Day 14 but distribute analysis steps over Days 14-18.) In our experience, airway organoids can be passaged at least 8 times (approximately 4 months), while alveolar organoids can be passaged 3-4 times; approximately 1 in 10 alveolar lines have survived to passage 8 or higher.

### Differentiating Airway Organoids at Air-Liquid Interface (ALI)

Airway organoids grown under the conditions above consist of >80% basal cells (KRT5+ TP63+ NGFR+) (ref. 14 and **Figure 1D**). While the maintenance of stem cells is desirable for long-term organoid expansion, native airways contain mostly differentiated cell types such as secretory (also called club), goblet, and ciliated cells, which are uncommon in the 3D organoid cultures. To mimic native airways more closely, air-liquid interface (ALI) culture is a well-established model system, in which large or small airway epithelial cells are grown on a 2D semi-permeable membrane, with the apical side exposed to air and the basolateral side exposed to media.^17,18^ Most methods involve expanding primary or commercial basal cell lines in traditional 2D culture, then transferring cells into a 24-well Transwell insert for ALI. These 2D-to-ALI methods require significant numbers of stem cells to establish the initial cultures and become less reliable at higher passages.^18^ A few groups have expanded cells as 3D organoids instead, then moved into ALI.^5,12,13,19,20^ We and others have shown that these 3D-to-ALI methods allow for the use of starting material that contains only rare stem cells—for example, BAL, tracheal aspirates, and nasal turbinate brush specimens. Here, we expand on this finding by establishing the first time course of differentiation from BAL-derived organoids to determine whether BAL-derived stem cells retain full differentiation potential.

We briefly describe our step-by-step protocol using commercially-available, serum-free PneumaCult-Ex Plus (basal cell expansion), PneumaCult-ALI (large airway differentiation), and PneumaCult-ALI-S (small airway differentiation) media. Media were prepared according to the manufacturer’s instructions and supplemented with Primocin antimicrobials to a final concentration of 100 µg/ml.

C) Approximate time for setting up ALI: 1 hr
  C1) Dissociate airway organoids (usually Passage 3) to a single-cell suspension as described in the passaging protocol, steps **B2**-**B12**.
  C2) Resuspend the cell pellet in Complete PneumaCult-Ex Plus Media to a concentration of 40,000 cells per 200 µl.
    - **Note:** We do not recommend using all cells for ALI, since ALI cultures cannot be passaged further. We set aside a portion of cells at P3 for ALI and either continue to expand the rest in 3D organoid culture or cryopreserve the excess cells.
  C3) Add 200 µl of cell suspension to the apical (top) side of a Transwell insert in a 24-well plate, for as many wells as desired. Add 500 µl of Complete PneumaCult-Ex Plus Media (without cells) to the basolateral (bottom) side. Incubate cells at 37°C, 5% CO2.
  C4) Feed with Complete PneumaCult-Ex Plus Media to both apical and basolateral sides every 2-3 days until cells are 90-100% confluent, usually 3 days.
  C5) Gently aspirate media from the apical side, being careful not to touch the cells. On the basolateral side only, replace PneumaCult-Ex Plus with 500 µl PneumaCult-ALI or ALI-S, leaving the apical side of cells exposed to air. This is termed “air lift” and is Passage 3 Day 0 of ALI.
    - *Tip:* Lift the inserts with sterile forceps and tilt them to aspirate while keeping the pipette tip away from the cell layer. Stabilize the inserts so that they do not move when aspirating or pipetting.
  C6) After air lift, change media on the basolateral side every Monday/Wednesday/Friday (2-3 days).
  C7) Once per week, add 200 µl of pre-warmed DPBS to the apical side and gently aspirate to wash away excess mucus.
  C8) Fix ALI at desired time points: Transfer the Transwell insert to a new 24-well plate; do not allow cells to dry out. Wash both apical and basolateral sides with DPBS (200 µl on top, 500 µl on bottom), add 10% formalin to both sides, and incubate at room temperature for 30 min. Wash with DPBS, then fill the well with 2 ml DPBS to store the fixed Transwell inserts. Store at 4°C.
  C9) Quantify the major airway cell types using immunofluorescence (described below).

#### ALI: Protocol Variations

- Instead of dissociating airway organoids and plating directly into ALI culture, the single-cell suspension can be sorted using flow cytometry for NGFR+ basal cells, as previously described.^14^
- A variation of airway organoid media has been developed that promotes differentiation into ciliated cells in 3D culture, though secretory and goblet cells were lost.^13^ Thus, 2D ALI remains the method of choice to recapitulate the diversity of differentiated airway cell types, but the 3D method offers an alternative model for ciliary diseases.

### Differentiating Alveolar Organoids in 3D Culture

Similarly, we aimed to determine whether BAL-derived alveolar organoids, which contain >80% alveolar type 2 cells (SFTPC+ HT2-280+) (**Figure 1D**), could be differentiated into alveolar type 1 (AT1) cells. Traditionally, alveolar progenitor cells have been grown in 2D culture, on plastic or cover slips, in serum-containing media, which induces differentiation to AT1 cells.^21^ More recently, it was shown that alveolar organoids could be kept in 3D culture but switched into media containing 10% human serum (named Alveolar Differentiation Media, ADM) to produce AT1 cells in the organoids.^6^ Another group reported the first serum-free method to differentiate alveolar organoids derived from iPSCs, likewise keeping organoids in 3D culture and switching the media.^22^ The serum-free media was called L-DCI for its key components, with “L” denoting LATS inhibitor, which induces nuclear activation of YAP/TAZ; this was shown to be sufficient for AT2-to-AT1 differentiation.

To test whether our BAL-derived AT2 organoids could differentiate into AT1 cells, especially using a serum-free method, we grew Passage 3 alveolar organoids for 7 days in our Alveolar Media, then transitioned into ADM or L-DCI for another 7 days. As a control, we treated lung tissue-derived organoids in the same manner. At Passage 3 Day 14, organoids were fixed and stained as described in the next section.

### Characterizing Intact Organoids using Immunofluorescence

After experimenting with many different methods of fixing and staining organoids, we have found the following protocols to be optimal for maintaining organoids’ intact structure while removing Matrigel. Antibody dilutions that we have tested are included in **Table 1**, and the antibody panels used in the figures are given in **Table 3**.

**Table 3.** Antibody panels used with these protocols.

| Primary antibodies | Primary Ab dilution | Secondary antibodies | Secondary Ab dilution |
| --- | --- | --- | --- |
| Airway Differentiation |  |  |  |
| Chicken Polyclonal IgY anti-Human KRT5 (BioLegend 905901)* | 1:500 | Donkey anti-Chicken IgY Highly Cross Adsorbed, Alexa Fluor 647 (Thermo A78952) | 1:500 |
| Rabbit Monoclonal IgG anti-Human Acetyl-α-Tubulin (Cell Signaling 5335S) | 1:200 | Donkey anti-Rabbit IgG Highly Cross-Adsorbed, Alexa Fluor Plus 488 (Thermo A32790) | 1:500 |
| SCGB1A1 panel: Mouse Monoclonal IgG1k anti-Human SCGB1A1 (Santa Cruz sc-365992) | 1:200 | Donkey anti-Mouse IgG Highly Cross-Adsorbed, Alexa Fluor Plus 594 (Thermo A32744) | 1:500 |
| MUC5AC panel: Mouse Monoclonal IgG1k anti-Human MUC5AC (Thermo MA5-12178) | 1:200 |  |  |
| *As the chicken anti-KRT5 antibody has become unavailable, we tested an alternative panel: |  |  |  |
| Goat Monoclonal IgG anti-Human KRT5 (Abcam ab321868) | 1:500 | Donkey anti-Goat IgG Highly Cross-Adsorbed, Alexa Fluor Plus 594 (Thermo Fisher A32758) | 1:500 |
| Rabbit Monoclonal IgG anti-Human Acetyl-α-Tubulin (Cell Signaling 5335S) | 1:200 | Donkey anti-Rabbit IgG Highly Cross-Adsorbed, Alexa Fluor Plus 488 (Thermo A32790) | 1:500 |
| SCGB1A1 panel: Mouse Monoclonal IgG1k anti-Human SCGB1A1 (Santa Cruz sc-365992) | 1:200 | Donkey anti-Mouse IgG Highly Cross-Adsorbed, Alexa Fluor Plus 647 (Thermo A32787) | 1:500 |
| MUC5AC panel: Mouse Monoclonal IgG1k anti-Human MUC5AC (Thermo MA5-12178) | 1:200 |  |  |
| Alveolar Differentiation |  |  |  |
| Rabbit Polyclonal IgG anti-Human pro-SPC (Seven Hills WRAB-9337) | 1:500 | Donkey anti-Rabbit IgG Highly Cross-Adsorbed, Alexa Fluor Plus 488 (Thermo A32790) | 1:500 |
| Goat IgG anti-Human AGER (R&D Systems AF1145) | 1:200 | Donkey anti-Goat IgG Highly Cross-Adsorbed Secondary Antibody, Alexa Fluor Plus 594 (Thermo A32758) | 1:500 |

- **Important:** For steps involving transfer of large organoids, use a 1000-µl pipette with about 3 mm cut off to widen the tip. Cut tips using a sterile blade in a sterile Petri dish.

D) Approximate time for fixing: 2-3 hr
  D1) For each line of organoids to be fixed, coat all sides of a 5- or 15-ml tube with 1 ml 1% BSA in DPBS, which helps prevent cells from sticking to the plastic. Aspirate dry.
  D2) Aspirate culture media (save if desired) and wash the wells once with DPBS, without disturbing the Matrigel drop.
  D3) Add 1 ml of cold DPBS + 10 mM EDTA per well. While ejecting, use the pipette tip to scrape Matrigel gently off the bottom of the plate. Pipette up and down one time to break up Matrigel without disrupting the organoids. Transfer well contents to the conical tube. Wash well(s) with another 1 ml DPBS-EDTA to recover as many cells as possible.
    - **Note:** This step is similar to adding TrypLE when passaging organoids at step **B2**. Two wells of the same condition may be combined into one. The difference is that cold DPBS with EDTA will break up Matrigel without dissociating the cells.
  D4) Lay the tube on its side and gently agitate at 4°C for 1 hr to dissolve Matrigel. Make sure liquid is sloshing around.
  D5) Let tube sit upright for 10 min so that organoids sink by gravity. *Do not spin*, since spinning will crush the organoids’ structure. The organoids should form a loose pellet, visible to the naked eye.
  D6) Carefully aspirate DPBS-EDTA.
  D7) Add 1 ml 10% formalin *without touching the organoids*, which will become sticky. Use the flow of fluid out of the pipette to resuspend the organoids. Add another 1 ml formalin (total 2 ml).
  D8) Let sit for 30 min (maximum 2 hr) at room temperature to fix. Most organoids should sink.
  D9) Remove formalin to a designated waste container. Add 3 ml DPBS without touching the organoids.
  D10) Let sit for 10 min, aspirate, add another 3 ml DPBS and store at 4°C until ready for staining.

E) Approximate time for immunofluorescence day 1: 3 hr
  E1) Start with fixed organoids in DPBS.
    - **Important:** Allow organoids to sink by gravity; do not centrifuge. Some loss is inevitable, so start with extra organoids if possible.
  E2) Carefully aspirate old DPBS.
  E3) Resuspend organoids in 500 µl Permeabilization buffer (0.2% Triton X-100 in DPBS) and transfer to a labeled 48-well plate. Use extra buffer to collect as many organoids as possible. Let the organoids settle to the bottom of the 48-well plate for at least 2 min, then aspirate any excess volume, leaving approximately 700 µl of Permeabilization buffer per well.
  E4) Gently shake at room temperature for 15 minutes.
  E5) While waiting, prepare Blocking buffer (5% normal donkey serum + 0.1% Triton X-100 in DPBS). Calculate how much will be needed: 500 µl/well for blocking, 200 µl/well for primary antibody incubation, and 200 µl/well for secondary antibody incubation. Make enough for 1-2 extra wells to allow for pipetting error.
  E6) Aspirate Permeabilization buffer. *Always leave a cushion of 150-200 µl to avoid aspirating the organoids* (e.g. if starting with a total of 700 µl, remove no more than 550 µl).
    - *Tip:* To aspirate, use a 200-µl pipette. Remove bubbles first. Then, place the 200-µl tip just below the meniscus of the solution and follow it down while aspirating.
    - *Tip:* Pipette 200 µl of buffer into an empty well as a reference for how much volume to leave on top of the organoids.
    - *Tip:* Look inside the pipette tip to make sure no organoids were accidentally picked up. If they were, eject, let the organoids settle, and try again.
  E7) Add 500 µl Blocking buffer per well. Eject the buffer down the side of each well, without touching the well contents. This helps mix the organoids without having them stick to the pipette tip.
  E8) Gently shake at room temperature for 1 hour.
  E9) Make a solution containing all primary antibodies combined in Blocking buffer. Remember to make enough for 1-2 extra wells. Negative controls will get Blocking buffer only, or an isotype control if available.
    - **Note:** The 150-200 µl cushion will result in the primary antibody solution being diluted by up to 2-fold. Consider making the stock antibody solution at 2X the desired concentration, so that the final antibody incubation happens at the correct dilution. For example, make 1:250 stock to target 1:500 for the incubation.
  E10) Aspirate Blocking buffer. Add 200 µl of primary antibody solution to each well (*not* including negative controls). Add 200 µl of Blocking buffer to negative control wells.
  E11) Gently shake at 4°C overnight.

F) Approximate time for immunofluorescence day 2: 5-6 hr
  F1) Aspirate primary antibody solution, keeping the 150-200 µl cushion over the organoids at all times.
  F2) Add 500 µl Wash buffer (0.1% Triton X-100 in DPBS) down the side of each well, again using the flow of liquid to mix the organoids without touching them.
  F3) Gently shake at room temperature for 5 minutes. Aspirate Wash buffer.
  F4) Repeat steps **F2-F3** for a total of 5 washes.
    - **Note:** Given the volume left over each time, each wash is really a dilution, so it is important to complete 5 washes.
  F5) Make a solution containing all secondary antibodies combined in Blocking buffer. Remember to make enough for 1-2 extra wells. This time, include negative controls.
    - **Important:** Protect fluorescent antibodies from light by wrapping tubes and plates in aluminum foil.
  F6) Add 200 µl of secondary antibody solution to each well (*including* negative controls).
  F7) Gently shake at room temperature for 2 hr.
  F8) Aspirate secondary antibody solution.
  F9) Repeat steps **F2-F3** for a total of 5 washes. If desired, *add DAPI to the wash buffer for the fourth wash (final concentration 1 µg/ml)*.
  F10) Using 500 µl of DPBS, transfer the organoids from the 48-well plate to individual microfuge tubes. Use another 500 µl of DPBS to wash out the wells, collecting as many organoids as possible. Let the organoids sink to the bottom.
  F11) While waiting for organoids to settle, cut about 3 mm off of 200-µl pipettes to widen the tips. Label glass microscope slides and draw a circle (about 2 cm in diameter) on each slide with a hydrophobic marker.
  F12) Aspirate as much supernatant as possible without losing organoids.
  F13) Using the cut-off tips, add 30 µl of Prolong Gold Mountant to each sample. Gently resuspend the organoids in mountant and transfer into the circle on the pre-labeled slide. Use the pipette tip to move the majority of organoids toward the middle of the circle. Aspirate bubbles if necessary.
  F14) Place a coverslip on each slide, pressing very gently to remove any bubbles.
  F15) Store slides in the dark at room temperature overnight to let the mountant dry. Image on a confocal microscope or other instrument as desired, using the same microscope settings and image analysis settings for all comparable samples.

#### Immunofluorescence: Protocol Variations

- We used a modified version of this protocol to stain ALI membranes for the differentiation time course. In this case, we stained for KRT5+ basal cells, SCGB1A1+ secretory cells, MUC5AC+ goblet cells, acetylated tubulin-positive (AcTub+) ciliated cells, plus F-actin to outline all cells. Due to the number of markers used, we separated the staining into two antibody panels (**Table 3**). After fixing the ALI cultures at each time point, we cut each membrane in half, then removed the halves from the Transwell using a razor blade. Each half was stained in a 48-well plate using the methods above. The main difference was that, since the cells were still attached to the membrane, all the volume could be aspirated with each wash, as long as the cells were not allowed to dry out. Instead of DAPI at step **F9**, phalloidin was added to stain F-actin, according to the manufacturer’s instructions; this made cell counting easier. Membranes were mounted onto slides with the cell side facing the cover slip (plastic membrane facing the slide). After imaging, cells were manually counted in an area of 25,000-100,000 µm^2^.
- If only one antibody panel is desired, ALI membranes can be stained directly in the Transwell, adding reagents to the apical and basolateral sides, then cutting out and mounting the membrane as a last step. ALI membranes can also be paraffin-embedded and stained by standard methods.
- This protocol can be further adapted to stain Cytospin slides or formalin-fixed, paraffin-embedded (FFPE) organoids. FFPE samples require dewaxing and antigen retrieval. We have found that similar dilutions of the antibodies described work well across these various applications. The only format that has not worked well in our hands is “whole mount” staining of organoids still embedded in Matrigel in the culture well, perhaps because it is harder for antibodies to penetrate the Matrigel.

## RESULTS AND DISCUSSION

We optimized and standardized a pipeline for establishing, expanding, and differentiating airway and alveolar organoids from human BAL fluid and showed that all major lung epithelial cell types can be derived from a single, accessible source of primary cells. We have intentionally designed all steps to be accessible to users with standard laboratory skills and resources. All media components are chemically-defined and serum-free, which improves reproducibility by reducing lot-to-lot variation and avoiding exposure to serum components that can change cellular identity. We were the first group to report BAL-derived alveolar organoids, but previously, the yield of AT2 cells ranged widely, from 3-93% of epithelial cells. We have now improved upon the technique so that we are consistently achieving >80% AT2 cells in our alveolar organoids.

After plating fresh BAL specimens, we counted and imaged organoids at Passage 0 Day 10 (**Figure 1C**). At this point, 96% of specimens (n = 45) yielded organoids in Airway Media, and 93% (n = 44) yielded organoids in Alveolar Media (updated from ref. 14). Organoids have variable morphologies initially but have a well-defined border that distinguishes them from the background of immune cells. Immune cells drop out of the cultures after passaging, since the media selects for epithelial cell outgrowth, and the 200*g* spins help to eliminate dead cells and debris. About half of the BAL organoids were passaged using our updated, standardized pipeline: we were able to expand 88% of airway organoid cultures (n = 24) and 91% of alveolar organoid cultures (n = 22) to at least Passage 2 for further characterization. By Passage 2 Day 14-18, airway organoids are spherical, a mixture of solid and cystic depending on organoid density and days in culture. Alveolar organoids are usually solid with slightly irregular borders; thin-walled spheres can also be observed. We used flow cytometry to validate the organoids’ cellular composition. All cultures yielded >95% epithelial cells, even though the starting BAL fluid contained >95% immune cells (not shown).^23,24^ The epithelial population consisted, on average, of 94 ± 3% NGFR+ basal cells in airway organoids and 90 ± 8% HT2-280+ AT2 cells in alveolar organoids (**Figure 1D**). Details of our flow cytometry methods are given in ref. 14 and summarized in **Supplemental Figure S1**.

We speculate that the improvement of alveolar cell selection compared to our prior report stems from standardization of BAL procedures. Though we cannot test this directly, the volume of BAL is known to influence the composition of cells in the return fluid. The first 50-60 ml aliquot generally samples small airway cells, while the second aliquot reaches alveolar spaces.^25^ Since almost all patients at our center receive 100-120 ml of saline instillation, and the return is pooled, we are likely capturing more alveolar cells, as well as improving normalization across specimens.

The prevalence of airway and alveolar stem cells in the organoid cultures is highly desirable for continued expansion and studying cell proliferation phenotypes. However, to model the cellular diversity of the native airway or alveolar spaces, methods that promote differentiation into specific cell types are needed. We prioritized serum-free approaches. For differentiation of airway basal cells, 2D air-liquid interface culture remains the method of choice, using commercially-available serum-free media such as PneumaCult. In contrast, differentiation of AT2 cells traditionally requires serum in 2D methods, so we explored newer serum-free methods that maintain the 3D organoid format.

To demonstrate differentiation of airway basal cells, we transitioned BAL-derived airway organoids into 2D ALI cultures and generated a time course of differentiation into secretory, goblet, and ciliated cells (**Figure 2**). At Day 1 after air lift, the membranes were covered with flattened KRT5+ basal cells. We observed no difference when starting from unsorted airway organoid cells or NGFR+ sorted basal cells (not shown). This suggests that any non-basal cell types in the organoids do not adhere or grow in the initial ALI conditions, so airway organoids can be transitioned directly to ALI without sorting. Interestingly, a few bundles of AcTub were visible, corresponding to active mitosis and midbody formation, which is required for primary ciliary development (**Supplemental Figure S2**).^26,27^ By Day 7, SCGB1A1+ secretory cells, MUC5AC+ goblet cells, and AcTub+ ciliated cells were all emerging; some of these cells were double positive for KRT5+, likely representing intermediate states. Secretory and goblet cells peaked in number at approximately Day 14 though overall made up <10% of the total cell population. Ciliated cells predominated by Day 21, rising to approximately 40-60% of the total, and were slightly more prevalent in PneumaCult-ALI (large airway) compared to ALI-S (small airway) media, as expected. One sample was grown to Day 28, which showed further ciliary maturation, with an increase in AcTub+ cells and decrease in KRT5+/AcTub+ double positive cells. The time course of differentiation of BAL-derived organoids was similar to that for tissue-derived organoid controls (**Supplemental Figure S2**).

**Figure 2.**
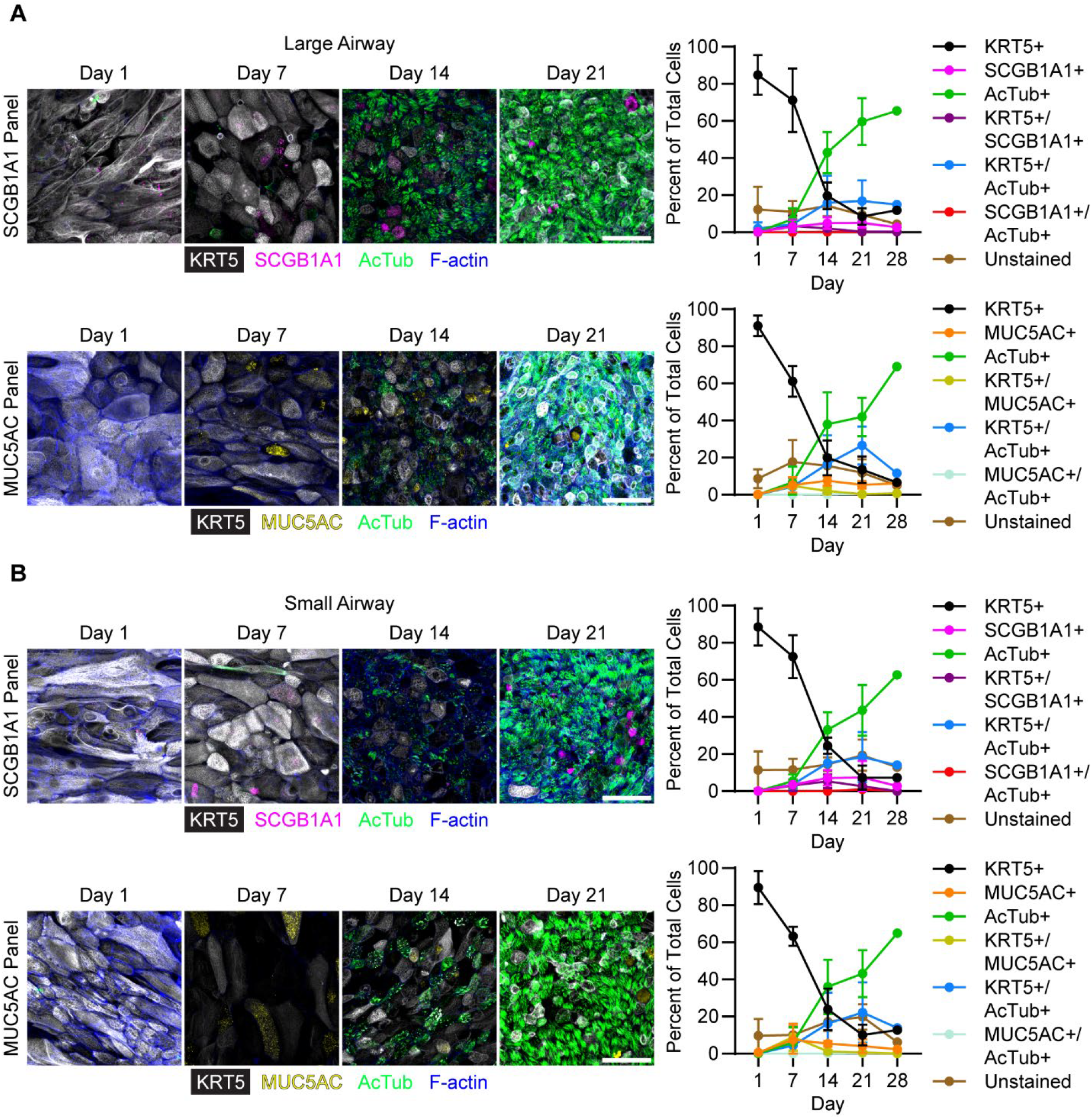
Time course of differentiation of BAL-derived airway organoids after being transitioned to air-liquid interface. Passage 3 cells grown in **(A)** PneumaCult ALI (large airway) media or **(B)** PneumaCult ALI-S (small airway) media were fixed at Day 1, 7, 14, and 21, then stained using two antibody panels for KRT5 (white), SCGB1A1 (magenta), MUC5AC (yellow), acetylated tubulin (AcTub, green), and F-actin (blue). Images were acquired on a Nikon A1R confocal microscope at 40X magnification with 0.5-µm slices. Each row of representative images was acquired and analyzed using the same settings and shown as maximum intensity projections. Scale bars are 50 µm. Cells were manually quantified and graphed as the mean ± SD from 5 BAL specimens. One specimen was extended to Day 28 for reference.

The cell proportions we observe at ALI Day 21-28 align well with the Human Lung Cell Atlas of primary airway tissue, suggesting that the BAL organoid-to-ALI approach can mimic native airways.^28,29^ However, our cell counts were limited by having to use two separate staining panels to accommodate five markers. We also did not stain for rare cell types such as respiratory airway secretory cells, ionocytes, etc., but estimate they would total <10% of all cells. Despite these limitations, we show overall that BAL-derived airway organoids can be differentiated as expected at ALI, and we identify the specific time points at which secretory, goblet, and ciliated cells are enriched.

To demonstrate differentiation of AT2 cells, we maintained BAL-derived alveolar organoids in 3D culture while modifying the media conditions. We tested two recently-published media formulations: serum-containing ADM^6^ and serum-free L-DCI^22^. As is typical for human alveolar organoids,^4,6,7,30^ we rarely observed AT1 cells in maintenance Alveolar Media, and the organoid morphologies ranged from thin-walled hollow spheres to more solid, irregular, and lobulated in appearance (**Figure 3A**). After 7 days in ADM or L-DCI differentiation media, organoids exhibited only subtle morphologic changes: some developed flatter patches or small budding cells, while others appeared unchanged. As expected, there was heterogeneity within and between specimens. Differences were more apparent after organoids were fixed and stained for the AT2 cell marker surfactant protein C (SFTPC) and the AT1 cell marker advanced glycosylation end-product specific receptor (AGER) (**Figure 3B**). In L-DCI media, 76% of BAL-derived organoids (n = 71) contained a majority of AGER+ cells, with a corresponding decrease in SFTPC signal. ADM had a less pronounced and more variable effect, with only 32% of organoids (n = 38) showing majority AGER expression. The extent of differentiation for BAL-derived organoids was the same or better than for tissue-derived controls. It is possible that differentiation in ADM would have been more extensive if allowed to continue for 10-14 days, but we stopped after 7 days to prevent cultures from becoming overgrown. Overall, we demonstrate proof of principle that BAL-derived organoids can respond to established, serum-free differentiation cues, such as air-liquid interface culture for airway cells and YAP/TAZ activation for alveolar cells.

**Figure 3.**
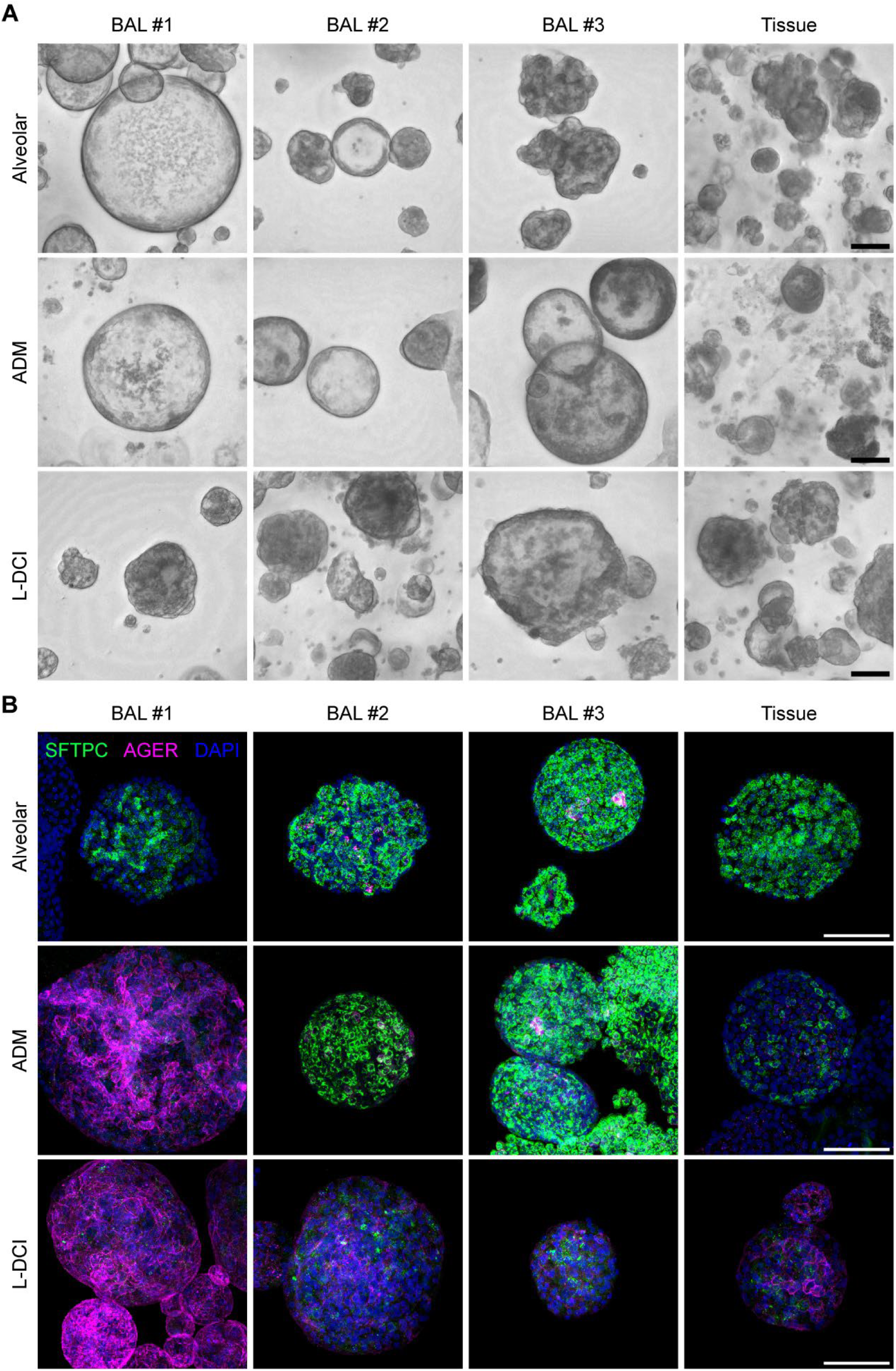
Differentiation of BAL-derived alveolar organoids. Passage 3 organoids were grown for 7 days in Alveolar Media, then switched into serum-containing (ADM, ref. 6) or serum-free (L-DCI, ref. 22) differentiation media for another 7 days, or kept in Alveolar Media. Tissue-derived organoids were used as controls. **(A)** Brightfield images of organoids at Day 14 were acquired on a Keyence BZ-X810 microscope at 20X magnification. The Keyence “full focus” feature was used to show multiple representative organoids for three BAL-derived cell lines and one tissue-derived control. **(B)** Organoids were freed from Matrigel, fixed intact, and stained for the AT2 cell marker SFTPC (green) and the AT1 cell marker AGER (magenta). Images were acquired on a Nikon A1R confocal microscope at 40X magnification with 2-µm slices, using the same settings throughout. Shown are maximum intensity projections of representative organoids. Scale bars are 100 µm.

## SUMMARY AND CONCLUSIONS

In summary, our results demonstrate that rare epithelial cells shed into BAL fluid can form organoids that behave in all the ways expected of organoids grown from lung tissue or iPSCs, from passaging to differentiation. By integrating organoid technology with BAL, a common clinical procedure, we highlight the opportunity to obtain primary lung cells for research with minimal risk to patients. Importantly, this protocol is readily adapted to other starting material; we have used cryopreserved human BAL, murine BAL, and human and murine lung tissue. We have not tried alternative airway specimens such as bronchial brushings, tracheal aspirates, nasopharyngeal swabs, or transbronchial biopsies. We speculate that these might be feasible as well, since the number of airway epithelial cells in such specimens is expected to be higher than in BAL fluid.^24,31^ Indeed, recent advances in the organoid field highlight the model’s versatility and its remarkable ability to generate organoids from unexpected sources of primary cells.^32^ However, it remains critical to validate whether organoids recapitulate the systems or diseases they are intended to model. We are currently investigating how BAL-derived organoid phenotypes correlate with disease processes such as lung transplant rejection.

While our models are so far limited to epithelial monocultures, they provide a platform to add further layers of complexity. Organoid and ALI cultures can both be exposed to chemicals, chemokines, or aerosols for mechanistic manipulations or drug discovery.^12,33–35^ Matrix material can be altered to examine the effects of extracellular proteins and mechanical tension.^36,37^ Epithelial cells can be co-cultured with immune, mesenchymal, or endothelial cells and infected with pathogens.^38–41^ Increasingly, cells from organoids are being altered genetically or transplanted.^10,42–45^ We envision that the combination of accessible starting material and straightforward protocols may facilitate these advanced applications.

## AUTHOR CONTRIBUTIONS

CFK and MYL conceived and designed research; TBG, BC, and MYL developed the protocols; TBG, BC, MSJ, AKK, MP, and MYL performed experiments; TBG, BC, SRR, MSJ, and MYL analyzed data; TBG, LMS, CFK, and MYL interpreted results; TBG, SRR, MSJ, AKK, SCP, and MYL prepared tables and figures; LMS, CFK, and MYL secured funding; MYL drafted the manuscript; and all authors edited and approved the manuscript.

## ACKNOWLEDGMENTS

We thank the many patients, physicians, nurses, and staff who helped provide BAL and tissue specimens at UW-Madison and Mass General Brigham. We greatly appreciated advice from Drs. Ruobing Wang and Stuart Rollins regarding ALI and Daniel Zeve regarding IF on intact organoids. We are grateful to the University of Wisconsin Optical Imaging Core and Carbone Cancer Center Flow Cytometry Laboratory (supported by P30 CA014520 and 1S10RR025483-01) for use of their instruments and services. Finally, we thank all members of the Schnapp and Kim labs for their collaboration and feedback.

## REFERENCES

1. Katsura, H. & Hogan, B. L. M. Lung organoids: powerful tools for studying lung stem cells and diseases. In Lung Stem Cells in Development, Health and Disease 175–189 (European Respiratory Society, 2021). doi:10.1183/2312508X.10009920.

2. Vazquez-Armendariz, A. I. & Tata, P. R. Recent advances in lung organoid development and applications in disease modeling. J. Clin. Invest. 133, (2023).

3. Rock, J. R. et al. Basal cells as stem cells of the mouse trachea and human airway epithelium. Proc. Natl. Acad. Sci. U. S. A. 106, 12771–12775 (2009).

4. Barkauskas, C. E. et al. Type 2 alveolar cells are stem cells in adult lung. J. Clin. Invest. 123, 3025–3036 (2013).

5. Sachs, N. et al. Long-term expanding human airway organoids for disease modeling. EMBO J. 38, 1–20 (2019).

6. Katsura, H. et al. Human Lung Stem Cell-Based Alveolospheres Provide Insights into SARS-CoV-2-Mediated Interferon Responses and Pneumocyte Dysfunction. Cell Stem Cell 27, 890-904.e8 (2020).

7. Youk, J. et al. Three-Dimensional Human Alveolar Stem Cell Culture Models Reveal Infection Response to SARS-CoV-2. Cell Stem Cell 27, 905-919.e10 (2020).

8. Wong, A. P. et al. Directed differentiation of human pluripotent stem cells into mature airway epithelia expressing functional CFTR protein. Nat. Biotechnol. 30, 876–882 (2012).

9. Huang, S. X. L. et al. Efficient generation of lung and airway epithelial cells from human pluripotent stem cells. Nat. Biotechnol. 32, 84–91 (2014).

10. Jacob, A. et al. Differentiation of Human Pluripotent Stem Cells into Functional Lung Alveolar Epithelial Cells. Cell Stem Cell 21, 472-488.e10 (2017).

11. Hawkins, F. J. et al. Derivation of Airway Basal Stem Cells from Human Pluripotent Stem Cells. Cell Stem Cell 28, 79-95.e8 (2021).

12. Eenjes, E. et al. Disease modeling following organoid-based expansion of airway epithelial cells. Am. J. Physiol.-Lung Cell. Mol. Physiol. 321, L775–L786 (2021).

13. van der Vaart, J. et al. Modelling of primary ciliary dyskinesia using patient-derived airway organoids. EMBO Rep. 22, e52058 (2021).

14. Liu, M. Y. et al. Human Airway and Alveolar Organoids from BAL Fluid. Am. J. Respir. Crit. Care Med. 209, 1501–1504 (2024).

15. Konishi, S., Tata, A. & Tata, P. R. Defined conditions for long-term expansion of murine and human alveolar epithelial stem cells in three-dimensional cultures. STAR Protoc. 3, 101447 (2022).

16. Martinu, T. et al. International Society for Heart and Lung Transplantation consensus statement for the standardization of bronchoalveolar lavage in lung transplantation. J. Heart Lung Transplant. 39, 1171–1190 (2020).

17. Fulcher, M. L., Gabriel, S., Burns, K. A., Yankaskas, J. R. & Randell, S. H. Well-Differentiated Human Airway Epithelial Cell Cultures. In Human Cell Culture Protocols (ed. Picot, J.) 183–206 (Humana Press, Totowa, NJ, 2005). doi:10.1385/1-59259-861-7:183.

18. Myo, Y. P. A. et al. Protocol for differentiating primary human small airway epithelial cells at the air-liquid interface. Am. J. Physiol.-Lung Cell. Mol. Physiol. 328, L757–L771 (2025).

19. Lee, W. et al. A single-cell atlas of in vitro multiculture systems uncovers the in vivo lineage trajectory and cell state in the human lung. Exp. Mol. Med. 55, 1831–1842 (2023).

20. Wang, R. et al. De Novo Generation of Pulmonary Ionocytes from Normal and Cystic Fibrosis Human Induced Pluripotent Stem Cells. Am. J. Respir. Crit. Care Med. 207, 1249–1253 (2023).

21. Dobbs, L. G. Isolation and culture of alveolar type II cells. Am. J. Physiol.-Lung Cell. Mol. Physiol. 258, L134–L147 (1990).

22. Burgess, C. L. et al. Generation of human alveolar epithelial type I cells from pluripotent stem cells. Cell Stem Cell 31, 657-675.e8 (2024).

23. Meyer, K. C. et al. An official American Thoracic Society clinical practice guideline: The clinical utility of bronchoalveolar lavage cellular analysis in interstitial lung disease. Am. J. Respir. Crit. Care Med. 185, 1004–1014 (2012).

24. Maestre-Batlle, D. et al. Novel flow cytometry approach to identify bronchial epithelial cells from healthy human airways. Sci. Rep. 7, 1–9 (2017).

25. Levy, L. et al. Sequential broncho-alveolar lavages reflect distinct pulmonary compartments: clinical and research implications in lung transplantation. Respir. Res. 19, 1–11 (2018).

26. Nekooki-Machida, Y. & Hagiwara, H. Role of tubulin acetylation in cellular functions and diseases. Med. Mol. Morphol. 53, 191–197 (2020).

27. Park, H.-S. et al. ARP-T1-associated Bazex–Dupré–Christol syndrome is an inherited basal cell cancer with ciliary defects characteristic of ciliopathies. Commun. Biol. 4, 544 (2021).

28. Travaglini, K. J. et al. A molecular cell atlas of the human lung from single-cell RNA sequencing. Nature 587, 619–625 (2020).

29. Sikkema, L. et al. An integrated cell atlas of the lung in health and disease. Nat. Med. 29, 1563–1577 (2023).

30. Alysandratos, K.-D. et al. Culture impact on the transcriptomic programs of primary and iPSC-derived human alveolar type 2 cells. JCI Insight 8, (2023).

31. Lu, J. et al. Rho/SMAD/mTOR triple inhibition enables long-term expansion of human neonatal tracheal aspirate-derived airway basal cell-like cells. Pediatr. Res. 89, 502–509 (2021).

32. Gerli, M. F. M. et al. Single-cell guided prenatal derivation of primary fetal epithelial organoids from human amniotic and tracheal fluids. Nat. Med. 30, 875–887 (2024).

33. Schamberger, A. C., Staab-Weijnitz, C. A., Mise-Racek, N. & Eickelberg, O. Cigarette smoke alters primary human bronchial epithelial cell differentiation at the air-liquid interface. Sci. Rep. 5, 8163 (2015).

34. Berical, A. et al. A multimodal iPSC platform for cystic fibrosis drug testing. Nat. Commun. 13, 4270 (2022).

35. Suezawa, T. et al. Disease modeling of pulmonary fibrosis using human pluripotent stem cell-derived alveolar organoids. Stem Cell Rep. 16, 2973–2987 (2021).

36. Shao, Z., De Coppi, P. & Michielin, F. Leveraging mechanobiology and biophysical cues in lung organoids for studying lung development and disease. Front. Chem. Eng. 5, (2023).

37. De Santis, M. M., Michielin, F., Shibuya, S., Coppi, P. de & Wagner, D. E. Lung tissue bioengineering for transplantation and modelling of development, disease and regeneration. In Lung Stem Cells in Development, Health and Disease 248–272 (European Respiratory Society, 2021). doi:10.1183/2312508X.10011020.

38. Tan, Q., Choi, K. M., Sicard, D. & Tschumperlin, D. J. Human airway organoid engineering as a step toward lung regeneration and disease modeling. Biomaterials 113, 118–132 (2017).

39. Dijkstra, K. K. et al. Generation of Tumor-Reactive T Cells by Co-culture of Peripheral Blood Lymphocytes and Tumor Organoids. Cell 174, 1586-1598.e12 (2018).

40. Yonker, L. M. et al. Development of a Primary Human Co-Culture Model of Inflamed Airway Mucosa. Sci. Rep. 7, 8182 (2017).

41. Ament, A.-L. et al. Endothelialized Bronchioalveolar Lung Organoids Model Endothelial Cell Responses to Injury. Am. J. Respir. Cell Mol. Biol. https://doi.org/10.1165/rcmb.2023-0373MA (2024) doi:10.1165/rcmb.2023-0373MA.

42. Ararat, E. et al. Comparison of Transplantation of Lung Organoid Cell Types: One Size Does Not Fit All. Am. J. Respir. Cell Mol. Biol. 66, 340–343 (2022).

43. Louie, S. M. et al. Progenitor potential of lung epithelial organoid cells in a transplantation model. Cell Rep. 39, 110662 (2022).

44. Ma, L. et al. Airway stem cell reconstitution by the transplantation of primary or pluripotent stem cell-derived basal cells. Cell Stem Cell 30, 1199-1216.e7 (2023).

45. Herriges, M. J. et al. Durable alveolar engraftment of PSC-derived lung epithelial cells into immunocompetent mice. Cell Stem Cell 30, 1217-1234.e7 (2023).

