## Supplemental Material for "Standardized pipeline for establishing, expanding, and differentiating airway and alveolar organoids from human BAL fluid"

**Supplemental Figure S1.** Summarized methods and gating strategies for flow cytometry.

**Supplemental Figure S2.** Differentiation of organoids at ALI.

**Supplemental Figure S1**

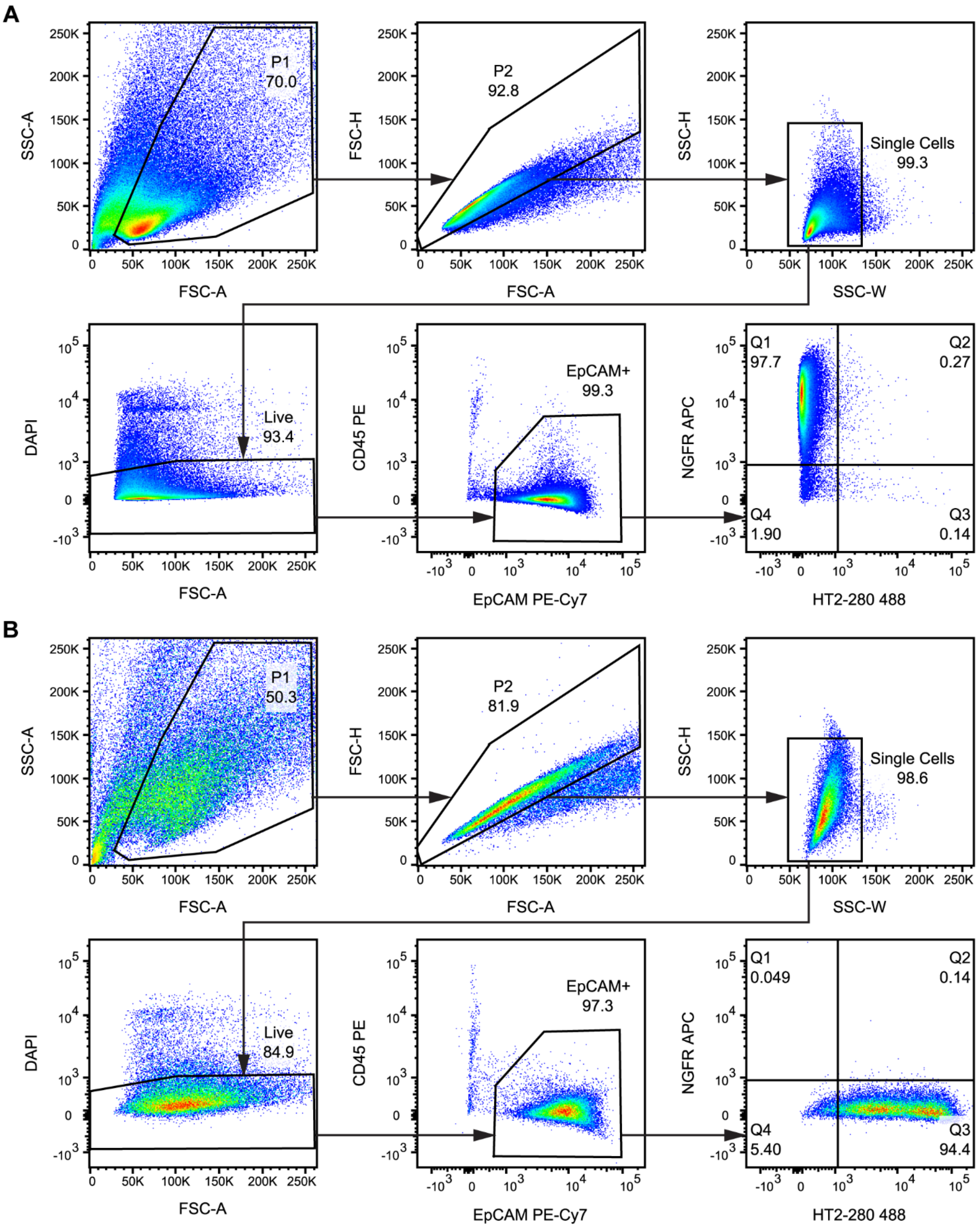

**Supplemental Figure S1**

Summarized methods and gating strategies for flow cytometry. Adapted from ref. 14. In brief, airway and alveolar organoids were dissociated to single cells at Passage 2 Day 14-18. Cells were passed through a 40-μm filter and collected into 100 μl of FACS Buffer (DPBS with 10% FBS, 2 mM EDTA, and 10 μM Y27632), with no more than 1 million cells per 100 μl. For controls, we used a mixture of all samples analyzed or compensation beads, as appropriate. Samples and controls were stained first with mouse IgM anti-human HT2-280 primary antibody at 1:50 dilution for 15 min on ice. To this, goat anti-mouse IgM Dylight 488 secondary antibody was added at 1:200 for 15 min. Samples were washed with FACS Buffer, resuspended again in 100 μl (per million cells), then stained using a combination of PE anti-human CD45, PE-Cy7 anti-human EpCAM, and APC anti-human NGFR fluorescent conjugated antibodies, each at 1:100 dilution for 15 min on ice. After washing again, cells were resuspended in 350 μl FACS Buffer and transferred to a 5-ml filter-cap tube for flow cytometry. DAPI was added to a final concentration of 2 μg/ml just prior to analysis on a BD FACS Aria II cell sorter. Representative results are shown for **(A)** airway and **(B)** alveolar organoids, which were analyzed in parallel with the same gating scheme. Q1 is the NGFR+ population and Q3 is the HT2-280+ population quantified in Figure 1D. Cells can be sorted into Passaging Buffer, washed, and plated into organoid or ALI culture. If replating into organoid cultures, we add human HRG1-β1 to Airway Media and human IL-1β to Alveolar Media to promote cell survival after sorting.

**Supplemental Figure S2**

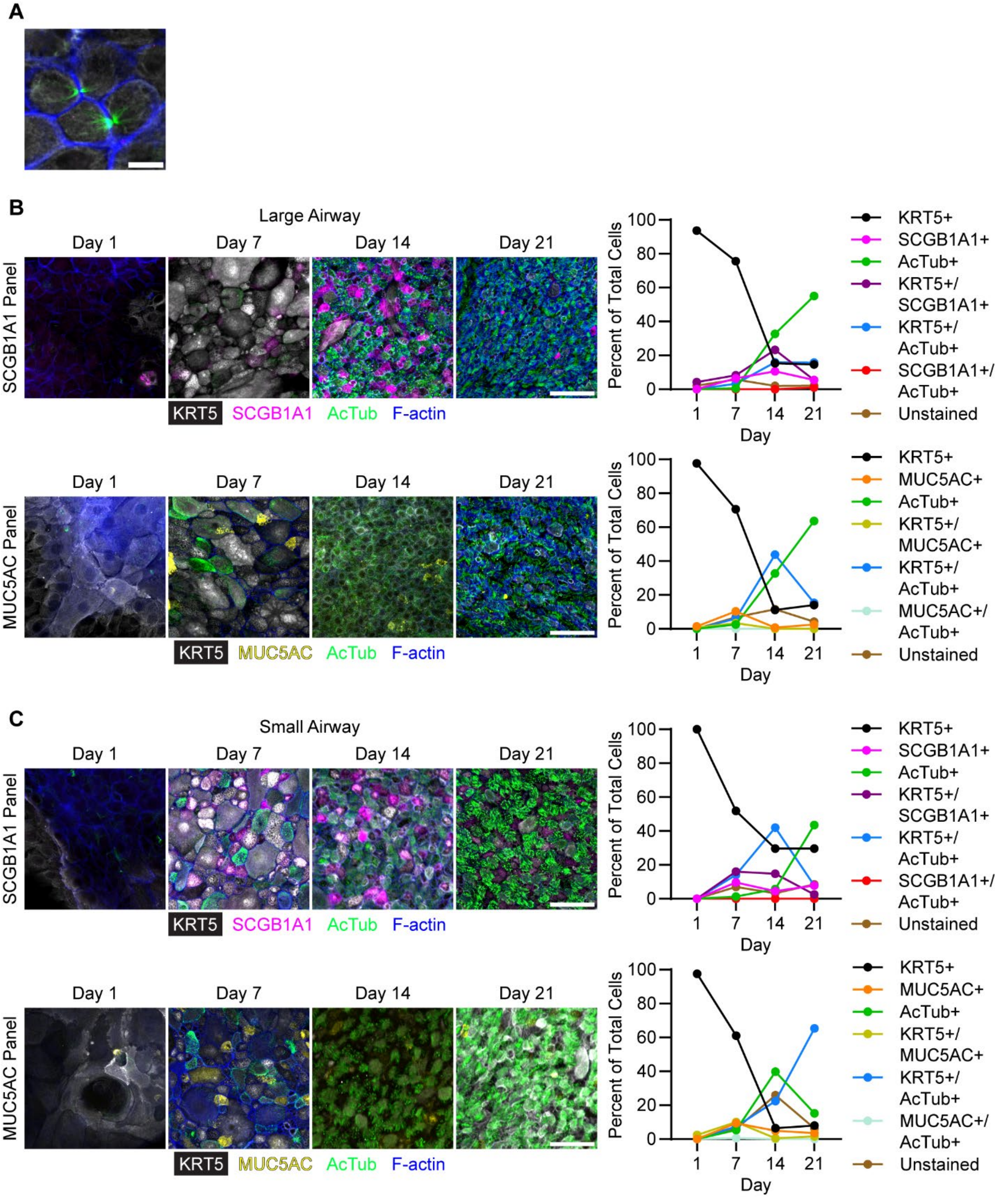

**Supplemental Figure S2**

Differentiation of organoids at ALI. **(A)** At Day 1, a few bundles of acetylated tubulin (green) are visible between adjacent KRT5+ (white) basal cells. These structures correspond to midbody formation during cytokinesis. Scale bar is 10 µm. As a control for differentiation, we used tissue-derived organoids transitioned to ALI in **(B)** PneumaCult ALI (large airway) media or **(C)** PneumaCult ALI-S (small airway) media. Manual cell counts were performed in the same manner as for Figure 2. Scale bars are 50 µm.
